# Multispecies translational evaluation of an NMDA receptor modulator reveals analgesic, autonomic-stabilizing, and stress mitigating effects without evidence of abuse liability

**DOI:** 10.64898/2026.09.14.751462

**Authors:** Joanne Tuohy, Toshitsugu Ishihara, Sudarshana Govindasamy, Ramu Anandakrishnan, Jennifer Davis, Fiona E. Harrison, Yuying Huang, Shao-Rui Chen, Hui-Lin Pan, Blaise M. Costa

## Abstract

CNS4 is an agonist concentration-biased NMDAR modulator exhibiting GluN2 subtype-dependent activity. Here we evaluated the translational pharmacology of CNS4 across mice, rats, and client-owned surgical oncology dogs. In mice, CNS4 did not produce conditioned place preference, indicating lack of reward liability. CNS4 produced dose-dependent reductions in locomotor activity without motor impairment on accelerating rotarod testing. CNS4 also maintained body temperature 0.5–1°C above the vehicle group in mice and dogs, consistent with NMDAR-dependent thermoregulatory activity and contrasting with hypothermic effects reported for NMDAR channel blockers such as ketamine. In fear-conditioning paradigms, CNS4 did not impair fear learning, memory consolidation, or fear expression. CNS4 reduced acute stress-induced sucrose preference, suggesting modulation of stress-responsive neural circuits. In a rat spinal nerve ligation model of neuropathic pain, both single-dose and three-day repeated CNS4 administration significantly reversed mechanical, pressure, and thermal hypersensitivity, supporting analgesic efficacy with no apparent evidence of rapid tolerance development. In client-owned surgical oncology dogs undergoing standard veterinary procedures, preliminary findings from 14 dogs (8 CNS4-treated, 6 vehicle-treated) demonstrated stable intraoperative mean arterial pressure and heart rate. Postoperatively, CNS4-treated dogs exhibited normal physiological recovery, with respiratory rate, heart rate, and body temperature returning toward the normal canine range, and required approximately 8% less propofol for anesthetic induction. Collectively, these findings support CNS4 as a novel NMDAR modulator with thermoregulatory, anti-agitative, analgesic, and autonomic-stabilizing properties without evidence of abuse liability, supporting continued translational development for neuropathic pain and possibly neuropsychiatric disorders.

## 1. Introduction

CNS4, 4-fluoro-N-[[2-(3-pyridinyl)-1-piperidinyl]-sulfanylidenemethyl]benzamide, is a small organic compound with drug-like properties (1). CNS4 is an anabasine-derived compound synthesized from the naturally occurring piperidinyl pyridine alkaloid anabasine, which is found in plants such as Nicotiana glauca (tree tobacco) (2).

Previous studies have demonstrated that CNS4 modulates N-methyl-D-aspartate receptor (NMDAR) subtypes, including GluN1/2A, GluN1/2B, GluN1/2AB, GluN1/2C, and GluN1/2D receptors (1, 3). At a concentration of 30 µM, CNS4 produced subtype-dependent changes in both glycine- and glutamate-mediated NMDAR currents. CNS4 increased glutamate potency at GluN1/2A and GluN1/2AB receptors, increased glycine potency at GluN1/2B and GluN1/2AB receptors, and increased glycine efficacy at GluN1/2C and GluN1/2D receptors (1, 3). In contrast, CNS4 reduced glycine efficacy at GluN1/2B and GluN1/2AB receptors and reduced glutamate efficacy at GluN1/2AB and GluN1/2C receptors, demonstrating a distinct subtype-dependent pharmacological profile across NMDAR subtypes. CNS4 showed no voltage-dependent effect; however, potentiation of GluN1/2A inward currents attenuated in Na⁺-free conditions, and it exhibited Ca^2+^-dependent blockade of inward currents at GluN1/2D receptors (3). CNS4 sensitized ambient agonists, producing transient reversible currents during CNS4 pre-application in GluN1/2A and GluN1/2AB receptors, with a larger effect in GluN1/2AB receptors (3). In cultured primary rat cortical, striatal, and cerebellar neurons, CNS4 potentiated NMDA-induced Ca²⁺ influx, with the strongest reported potentiation observed at 300 µM NMDA in cortical and striatal neurons, and a smaller but significant increase in cerebellar neurons (1).

CNS4 is highly brain-penetrant and it rapidly distributes into the central nervous system following systemic administration, reaching Tmax in 15 minutes following intraperitoneal injection in mice (4). Further, CNS4 increases thermal escape latency in a hot-plate assay, indicating analgesic activity. CNS4 also reduces stress-associated behaviors during fear conditioning, including decreased stress-induced defecation and hyperarousal responses (4). Importantly, CNS4 does not alter freezing behavior, suggesting preservation of fear memory formation and recall (4).

A recent translational study of CNS4 demonstrates favorable pharmacokinetic and safety characteristics in adult beagle dogs (5). Following subcutaneous administration, CNS4 exhibits rapid absorption (Tmax 1.5–1.7 h), an elimination half-life of approximately 6 – 7 hours and remains detectable in plasma for up to 24 h. Oral administration produces rapid systemic exposure (Tmax ∼1 h) and substantially higher plasma concentrations than subcutaneous dosing, confirming oral bioavailability (5). In a 14-day safety study, single subcutaneous doses of 5, 10 and 25 mg/kg produced no clinically significant changes in behavior, food consumption, body weight, hematology, or clinical chemistry parameters (5). These findings establish CNS4 as a well-tolerated, brain-penetrant NMDAR modulator with predictable pharmacokinetics in a large-animal species, supporting further translational development.

CNS4 is not chemically related to any known opioid alkaloids (1, 3). Although high affinity NMDAR channel blocker phencyclidine and its analog ketamine are abusive drugs (6), memantine, an FDA approved NMDAR antagonist for Alzheimer’s disease does not exhibit abuse liability (7). CNS4 is an allosteric modulator that is not capable of completely blocking NMDAR. CNS4 is neither a channel blocker nor a competitive antagonist (1, 3). However, since CNS4 is likely to alter glutamatergic neurotransmission, it is important to evaluate its potential effects on reward-related behaviors. Furthermore, NMDARs play critical roles in learning, memory, and synaptic plasticity, making assessment of abuse liability and cognitive function essential components of the preclinical safety and translational evaluation of CNS4. Given the abuse potential and cognitive effects reported for some NMDAR-targeting agents, we evaluated the effects of CNS4 across a range of behavioral paradigms.

While rodent studies provide important information on efficacy and mechanism of action, evaluation in client-owned surgical oncology dogs offers a clinically relevant translational model that more closely reflects the complexity of naturally occurring disease and perioperative pain. These studies enable assessment of CNS4 under real-world veterinary conditions, including concurrent anesthesia, analgesics, surgical stress, and postoperative recovery, while providing preliminary safety and pharmacodynamic data in a large-animal species. Furthermore, demonstration of efficacy and tolerability in client-owned dogs would strengthen the translational rationale for the development of CNS4 as a therapeutic for pain and stress-related disorders.

## 2. Methods

### 2.1 CNS4 Hydrocholoride salt development and synthesis

CNS4 free base (molecular weight 343.42 g/mol) was converted to its hydrochloride salt using a procedure developed in the Costa lab. Briefly, CNS4 free base was dissolved in anhydrous chloroform, and an equimolar amount of hydrochloric acid was added as a 4.0 M solution in dioxane under continuous stirring at 4° C. The reaction mixture was maintained under stirring until complete salt formation occurred, after which the product was precipitated by the addition of methyl tert-butyl ether. The precipitated CNS4 hydrochloride salt (CNS4·HCl; molecular weight ≈ 379.9 g/mol) was collected by filtration and dried at 37°C to constant weight. Typical isolated yields ranged from 85% to 95% of theoretical yield. CNS4·HCl was soluble in 20% (w/v) hydroxypropyl-β-cyclodextrin (HPβCD) prepared in double-distilled water, achieving concentrations up to 30 mg/mL without the need for organic cosolvents or PH adjustments.

Larger quantity of CNS4·HCl was synthesized in Roshel Laboratories, Bangalore. The identity and purity of the resulting salt were confirmed by liquid chromatography–mass spectrometry (LC-MS), nuclear magnetic resonance (NMR) spectroscopy, and high-performance liquid chromatography (HPLC), demonstrating >99% purity. The dried CNS4·HCl was stored under desiccated conditions until formulation and subsequent pharmacological studies. For simplicity, CNS4·HCl salt is referred as CNS4 hereafter. Since single dose rat neuropathic pain experiments were performed before the CNS4·HCl synthesis, original CNS4 base was used for this experiment.

### 2.2 Animals and treatments

Approximately 10 weeks old male C57Bl/6 mice were obtained from Charles River laboratories. These mice were housed on a 12:12 light/dark cycle for the duration of the experiment. Mice were pseudo-randomly allocated by cage to one of four treatment conditions with n=8-9 per group: Vehicle control (20% w/v hydroxypropyl-β-cyclodextrin in sterile water), CNS4 50 mg/kg, CNS4 100mg/kg, and positive control of cocaine hydrochloride 10 mg/kg or ketamine 15 mg/kg. All mouse behavioral studies were undertaken in the Vanderbilt Mouse Neurobehavioral Laboratory Core facility.

### 2.3 Conditioned Place Preference

Mice were placed in a 2-chamber shuttle box (MedAssociates) and allowed to move freely between compartments for 10 mins., to determine if a chamber preference existed (day 1). Days 2-5 mice were injected twice a day (am and pm), 15 minutes prior to the start of the trial. The mice were intraperitoneally injected with saline in the morning and confined to 1 side of the shuttle box. In afternoon of the same day, the mice were then injected with their assigned treatment and placed with only access to the other side of the shuttle box. Drug/side pairings were balanced across groups. After 4 total days of injections, mice were tested for place preference in the absence of saline or drug treatment. On day 6, the mice were placed in the shuttle box and allowed to move freely between both sides. In each 10 minutes session, the distance traveled in each compartment was recorded as a measure of drug response, and place preference was determined on the final trial by percent time spent in each chamber.

### 2.4 Locomotor activity

Exploratory locomotor activity was measured in an open field chamber measuring 27 x 27 cm (MedAssociates), over a 60-min. period. Infrared beams and detectors automatically recorded movement in the open field.

### 2.5 Rotarod

The ability to maintain balance on a rotating cylinder was measured with a standard rotarod apparatus. The apparatus consists of a cylinder (approximately 3 cm in diameter, covered with textured rubber) which rotates at speeds accelerating from 4 to 40 rpm over 5 minutes. Mice were placed on the rotarod, confined to a section of the cylinder approximately 6.0 cm long by Plexiglas dividers. The cylinder rotates 40 cm above an area into which mice may safely fall. Latency at which mice fell from the rotating cylinder was measured, as well as time to first rotation (spinning completely around the cylinder for one full rotation). Maximal trial length was 300 sec. Rotarod performance was assessed three times daily for 2 days.

### 2.6 Implantation of transducers and temperature recording

Mice were briefly anesthetized using isoflurane during implantation. Using a 15-gauge needle, containing the TP-500 temperature transponder, implants were injected subcutaneously at the back of the neck just above the shoulder blades. Temperatures were read using the BMDS reader from Avidity by holding the wand approximately 1 inch from the animal while it remained in the home cage and recording the value. Mice were permitted at least 24 hours to recover from transponder implantation prior to measurement of drug-induced temperature change since stress can alter temperature.

A baseline temperature of each mouse was taken before any injections were given. The mice were then given their designated IP injection, and temperature readings were taken at 10 min, 20 min, 30 min, 40 min, 50 min, 60 min, 1.5 hours, 2 hours, 2.5 hours, 3 hours. 3.5 hours, 4 hours, 4.5 hours, and 5 hours post injection. Mice remained in their home cage between measurements.

### 2.7 Fear conditioning

On the first day, mice were placed in an experimental chamber in which three footshocks of 0.5 mA and 2-s duration were delivered via the floor of metal bars over a period of 8-min. Each footshock was preceded by a 30-sec. tone of approximately 70 dB. Mice were then returned to the home cage. The following day, a retention test was administered wherein mice placed in the same experimental chamber, for 4 min., and then returned to their home cage. Approximately 1 hour later, mice were placed in a different environment using plastic inserts to alter the floor and walls (given a curved shape) with the addition of vanilla scent in the surrounding sound attenuating chamber. After 2 minutes in silence in the new environment, the 70-dB tone was presented for a duration of 2 minutes, mice were then returned to their home cage. In each of the three sessions, freezing behavior was observed and recorded automatically (MedAssociates).

### 2.8 Shock threshold analysis

This task was performed as a control measure for all tasks that involve administration of a footshock. An ascending series of mild footshocks was delivered through the grid floor of an experimental chamber, 1-sec. in duration, in intensities of 0.075, 0.1, 0.15, 0.2, 0.25, 0.3, 0.35, 0.4, 0.45, 0.5 mA, with an interstimulus interval of at least 30 sec. The intensity at which mice a) flinched (typically 0.1-0.2); b) ran or jumped (typically (0.15-0.25); and c) vocalized (typically 0.2-0.3 if at all) were recorded. The session was terminated immediately after the first time the mice vocalized or at 0.5 mA. Following threshold testing all mice were given a 2 second foot shock measuring 0.5 mA to invoke a stress response prior to sucrose preference testing.

### 2.9 Sucrose Preference

Mice were given two water bottles in their home cage one day prior to sucrose preference testing to habituate the mice to the setup. On the day of the test, the mice were individually housed and provided with two water bottles once again. One bottle contained water and the other bottle contained a 1% w/v sucrose solution. Both bottles were weighed before the mice were placed in the cage, and then again, the next day. The difference in weight was used to calculate water consumption, as well as to determine if there was a preference.

### 2.10 Rat spinal nerve ligation (SNL) model of neuropathic pain

Adult Sprague Dawley rats (8–10 weeks old, Envigo) were used to induce neuropathic pain as described previously (8). Briefly, rats were anesthetized with 2–3% isoflurane, and the left L5 and L6 spinal nerves were exposed and ligated with 6–0 silk sutures separately under a surgical microscope. Sham surgery (the same surgical procedure without nerve ligation) was performed as the control. CNS4 was injected through intraperitoneal route as 100mg/kg dose.

### 2.11 Nociceptive behavioral tests

Mechanical pressure thresholds were assessed using a Randall-Selitto paw pressure testing apparatus (#2500, IITC Life Science, Woodland Hills, CA). The hindpaw was gently secured using the clamp, and a steadily increasing force was applied to the plantar surface until a withdrawal response was elicited (9, 10). The force at the time of withdrawal was recorded as the pressure threshold.

Tactile withdrawal thresholds were measured using calibrated von Frey filaments. Animals were placed individually in transparent plastic chambers placed on an elevated mesh floor. Filaments were applied perpendicularly to the plantar surface of the hindpaw with sufficient force to bend the filament for 6 s. A brisk paw withdrawal or flinching response was considered a positive response. If no response occurred, a filament of greater force was applied; if a response occurred, the next filament of lower force was tested. Six consecutive applications including the first positive response were used to calculate the 50% withdrawal threshold using the up–down method (11, 12).

Heat withdrawal latency was evaluated using a plantar thermal testing apparatus (#390G, IITC Life Science), as described previously (9, 13). Animals were placed on a glass platform maintained at 30°C, and a mobile radiant heat stimulus was applied to the plantar surface of the hindpaw until the animal lifted or licked the paw. The latency to hindpaw withdrawal was recorded as the withdrawal latency.

### 2.12 Client own-surgical oncology dog enrollment

Client-owned dogs scheduled to undergo oncologic surgical procedures at the Virginia-Maryland College of Veterinary Medicine were prospectively enrolled following written informed owner consent. The study protocol was reviewed and approved by the Virginia Tech Institutional Animal Care and Use Committee (IACUC 25-014). CNS4 subcutaneous injection formulation was prepared aseptically in a certified Class II biological safety cabinet. The final formulation was sterilized by passage through a 0.2-µm membrane filter and collected into a sterile vial prior to administration. Dogs were assigned in a blinded manner whenever possible to receive either CNS4 or vehicle control in addition to the standard perioperative medications prescribed by the attending surgeon and anesthesiologist. Administration of CNS4 or vehicle did not replace or alter any aspect of the established standard-of-care anesthetic, analgesic, or surgical management. All clinical decisions regarding anesthesia, surgery, and postoperative care were made independently by the attending veterinary team. CNS4 or vehicle was administrated via subcutaneous injection as single 10mg/kg dose. For this part of the study, CNS4 free base was dissolved in a custom-made vehicle as previously published studies on research dogs in 30mg/ml concentration (5). Maximum of 5ml volume was injected at one site. For one patient, the dose was restricted to 40kg although it was heavier, since it was clinically diagnosed being overweight with poor health condition. Fourteen dogs of diverse breeds were enrolled, including mixed-breed dogs (n = 6), Labrador Retrievers (n = 2), Great Danes (n = 2), and one each of Pug, Catahoula Leopard mix, French Bulldog, and Maltese mix. Eight dogs received CNS4 (6 males, 2 females; mean age 8.4 ± 2.2 years), and six received placebo (4 males, 2 females; mean age 6.7 ± 3.0 years).

### 2.13 Sample collection and monitoring

Blood samples for pharmacokinetic (PK) analysis were collected at approximately 1, 4, and 8 hours after dosing. Vital signs including heart rate, mean arterial pressure (MAP), respiratory rate, and temperature were monitored throughout the surgery. Safety and pain assessments were made at the following time points: 2, 4 and 6 hours after extubation by blinded trained observers using the Short Form of the Glasgow Composite Measuring Pain Scale (SF-CMPS) scoring system (14, 15). With the SF-CMPS system, assessment categories were scored from 0-20. The consent was obtained to collect minimal volume of blood samples only for three time points. Therefore, CNS4 concentrations after 8-hour time points could not be calculated in the PK study.

### 2.14 Data analysis

Data containing repeated measurements across time, test session, or experimental condition were analyzed using two-way repeated-measures ANOVA, with treatment as the between-subject factor and time or experimental condition as the within-subject factor, as appropriate. When significant main effects or interactions were identified, appropriate multiple-comparisons tests (Šídák’s or Tukey’s, as indicated) were performed, with multiplicity-adjusted P values reported; P < 0.05 was considered statistically significant. For datasets containing missing observations, a mixed-effects model with restricted maximum likelihood was used in place of repeated-measures ANOVA, followed by appropriate multiple-comparisons testing.

## 3. Results

Mice were randomly assigned to one of three treatment groups for the conditioned place preference (CPP) study. The groups consisted of vehicle control (20% w/v HPβCD; n = 8), CNS4 50 mg/kg (n = 9), CNS4 100 mg/kg (n = 9), cocaine hydrochloride 10 mg/kg (n = 8). Cocaine was included as a positive control because of its established rewarding properties and ability to induce conditioned place preference in rodents. For all other behavioral assays, including locomotion, rotarod, fear conditioning, sucrose preference, nest building, and related stress-associated behavioral studies, low dose (15 mg/kg) ketamine hydrochloride was used as a positive control because subanesthetic doses have been shown to produce rapid antidepressant-, stress-modulating-, and neuroplasticity-related effects in rodents (16). All treatments were administered by intraperitoneal injection, and behavioral testing was performed according to the individual assay protocols. Investigators remained blind to treatment allocation, during data collection and analysis whenever feasible.

### 3.1 CNS4 does not cause conditioned place preference (CPP) in mice

In the CPP assay, all four groups displayed comparable baseline chamber preferences before starting the treatments, except for CNS4 at 50 mg/kg showed a modest increase in time spent in the right side chamber (57.72%; P = 0.0483), **Figure 1A**. On the final drug-free CPP test day, CNS4 did not produce a significant conditioned place preference or aversion and did not alter locomotor activity during the test (**Figure 1B&C**). For chamber duration, there was no significant effect of side (F (1,23) = 1.433, P = 0.2434), treatment (F (2,23) = 0.000, P > 0.9999), or side vs treatment interaction (F (2,23) = 0.9103, P = 0.4164). Similarly, distance traveled was comparable between chambers and treatment groups, with no significant effect of chamber side (F (1,23) = 0.8291, P = 0.3720), treatment (F (2,23) = 2.554, P = 0.0996), or side vs treatment interaction (F (2,23) = 0.2011, P = 0.8193). Together, these findings indicate that repeated CNS4 conditioning at 50 or 100 mg/kg produced neither significant place preference nor place aversion, without evidence of altered locomotor activity during the drug-free CPP test.

**Figure 1.**
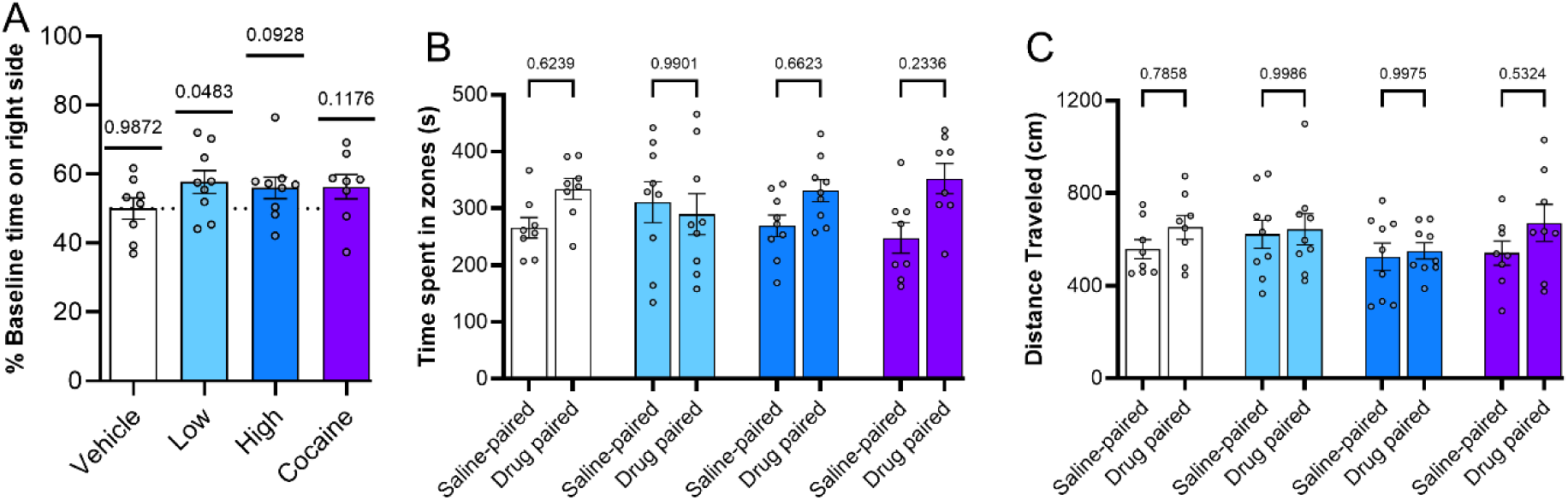
Conditioned place preference (CPP) assessment of CNS4 in mice. Male C57BL/6 mice were tested in a two-chamber (saline and treatment) shuttle box using a standard CPP paradigm. Following baseline preference assessment (A), mice underwent four days of alternating saline and treatment-paired conditioning sessions with balanced chamber assignments. On the final test day, mice were allowed free access to both chambers in the absence of drug treatment, and time spent in each chamber (B) and distance traveled (C) in each chamber were recorded. Groups included vehicle (20% HPβCD), no fill, n=8; CNS4 50 mg/kg (light blue), n=9; CNS4 and 100 mg/kg (dark blue), n=9; Cocaine 10 mg/kg (purple), n=8 administered intraperitoneally. Statistical analysis was performed using one sample t test to determine 50% probably of the baseline (before treatment) time spent in the right-side chamber (A); and two-way repeated-measures ANOVA followed by Šidák’s multiple-comparison test for the final test day comparisons presented in B&C.

### 3.2 CNS4 produces dose dependent locomotion effect without causing motor deficits

Exploratory locomotor activity was assessed in an open-field chamber for 60 min following drug administration. Two-way repeated-measures ANOVA revealed significant effects of treatment (F (3,30) = 30.45, P < 0.0001), time (F (11,330) = 100.8, P < 0.0001), and a significant treatment vs time interaction (F (33,330) = 9.856, P < 0.0001), indicating that the effects of treatment on locomotor activity varied over time (**Figure 2A**). Mice treated with CNS4 at 100 mg/kg exhibited markedly reduced locomotor activity compared with vehicle-treated animals during the early portion of the session. Significant reductions were observed at 5 min (mean difference = −468.1, P < 0.0001), 10 min (−535.5, P < 0.0001), 15 min (−508.1, P < 0.0001), 20 min (−412.3, P < 0.0001), 25 min (−355.8, P = 0.0001), 30 min (−333.8, P = 0.0004), 35 min (−233.2, P = 0.0302), 40 min (−229.4, P = 0.0347), 45 min (−282.2, P = 0.0043), and 50 min (−274.9, P = 0.0059).

**Figure 2.**
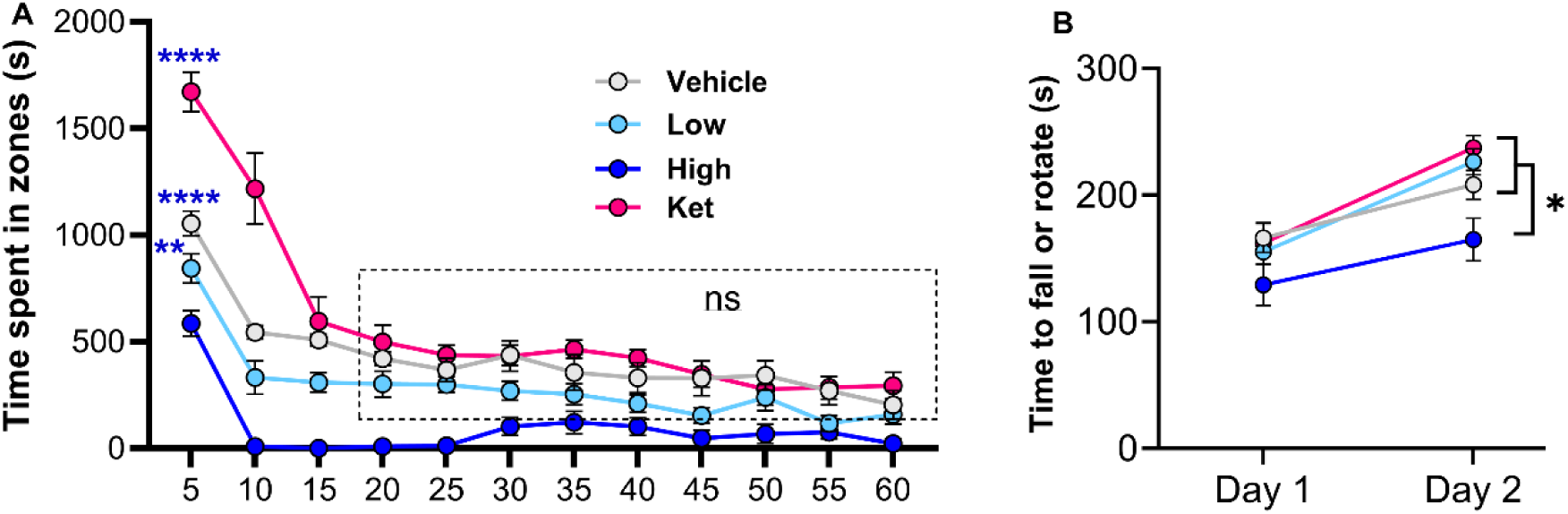
Effects of CNS4 on exploratory locomotor activity and motor coordination. Open-field locomotor activity was measured in automated infrared beam-equipped chambers over a 60-min period to assess exploratory behavior (A). Motor coordination and balance were evaluated using an accelerating rotarod apparatus with speeds increasing from 4 to 40 rpm over 5 min (B). Latency to fall and rotation (spinning completely around the cylinder for one full rotation) were automatically recorded. Blue asterisks represent statistical significance compared to high dose of CNS4. Dot box indicates the duration in which ketamine and CNS4 50mg/kg group were not different from vehicle. Route, intraperitoneal. *p<0.05, **p<0.01, ****p<0.0001.

Activity gradually recovered thereafter, with no significant differences from vehicle observed at 55 or 60 min. CNS4 at 100 mg/kg also significantly reduced locomotor activity relative to ketamine during most time bins and relative to CNS4 50 mg/kg during the first 25 min of testing. In contrast, CNS4 at 50 mg/kg did not differ significantly from vehicle at any time point (all P > 0.05). These findings demonstrate a dose-dependent suppression of exploratory locomotor activity following 100 mg/kg CNS4 administration, with the effect being most pronounced during the first 30–50 min after administration and largely resolving by the end of the observation period.

As expected, being a psychostimulant, ketamine produced a transient increase in exploratory locomotor activity during the early phase of testing. Ketamine-treated mice exhibited significantly greater activity than vehicle-treated mice at 5 min (mean difference = 617.3, P < 0.0001) and 10 min (mean difference = 673.4, P < 0.0001), with activity levels returning to those of vehicle-treated animals by 15 min and thereafter (all P > 0.05).

Speed escalating rotarod performance improved significantly from Day 1 to Day 2 across all treatment groups (F (1,30) = 40.77, P < 0.0001), demonstrating intact motor learning and task acquisition (**Figure 2B**). No significant day vs treatment interaction was observed (F (3,30) = 1.312, P = 0.2888), indicating that the rate of improvement was similar among groups. A significant main effect of treatment was identified (F (3,30) = 5.840, P = 0.0029), driven by a modest reduction in performance in mice treated with CNS4 100 mg/kg. Post hoc analyses revealed that the 100 mg/kg group exhibited lower latencies to fall compared to vehicle-treated mice (147.0 vs. 187.2 s, P = 0.0416), CNS4 50 mg/kg-treated mice (190.8 s, P = 0.0171), and ketamine-treated mice (199.7 s, P = 0.0041). In contrast, no differences were observed among the vehicle, CNS4 50 mg/kg, and ketamine groups. These findings suggest that CNS4 at 100 mg/kg produces a substantial reduction in exploratory behavior while causing only a mild reduction in rotarod performance and no detectable deficit in motor learning.

### 3.3 Dose dependent bidirectional effects of CNS4 on thermoregulation

Body temperature was monitored for 5 h following treatment using implanted subcutaneous transponders. Mixed-effects analysis revealed a significant effect of time (F (14,418) = 33.93, P < 0.0001) and a significant time vs treatment interaction (F (42,418) = 7.627, P < 0.0001), whereas the overall treatment effect was not significant (F (3,30) = 0.562, P = 0.6442), **Figure 3**. CNS4 at 100 mg/kg produced a distinct biphasic alteration in body temperature characterized by an early reduction relative to vehicle during the first 30 min after dosing, followed by a delayed increase above vehicle values beginning approximately 2.5 h post-dose. Compared with vehicle, body temperature in the CNS4 100 mg/kg group was significantly lower at 10 min (−1.36°C, P = 0.0003), 20 min (−1.46°C, P < 0.0001), and 30 min (−1.03°C, P = 0.0112). Subsequently, temperatures in the CNS4 100 mg/kg group exceeded those of vehicle-treated mice at 2.5 h (+1.13°C, P = 0.0041), 3 h (+1.08°C, P = 0.0071), 3.5 h (+1.01°C, P = 0.0145), 4 h (+1.02°C, P = 0.0130), 4.5 h (+1.08°C, P = 0.0087), and 5 h (+1.12°C, P = 0.0048). In contrast, neither CNS4 50 mg/kg nor ketamine differed significantly from vehicle at any time point. Although CNS4 100 mg/kg significantly altered the temporal profile of body temperature, the observed changes remained within the physiological range typically reported for mice (17) and therefore do not represent overt hypothermia or hyperthermia. Notably, the early decrease in body temperature coincided temporally with the transient suppression of exploratory locomotor activity observed in the open-field assay (**Figure 2A**).

**Figure 3.**
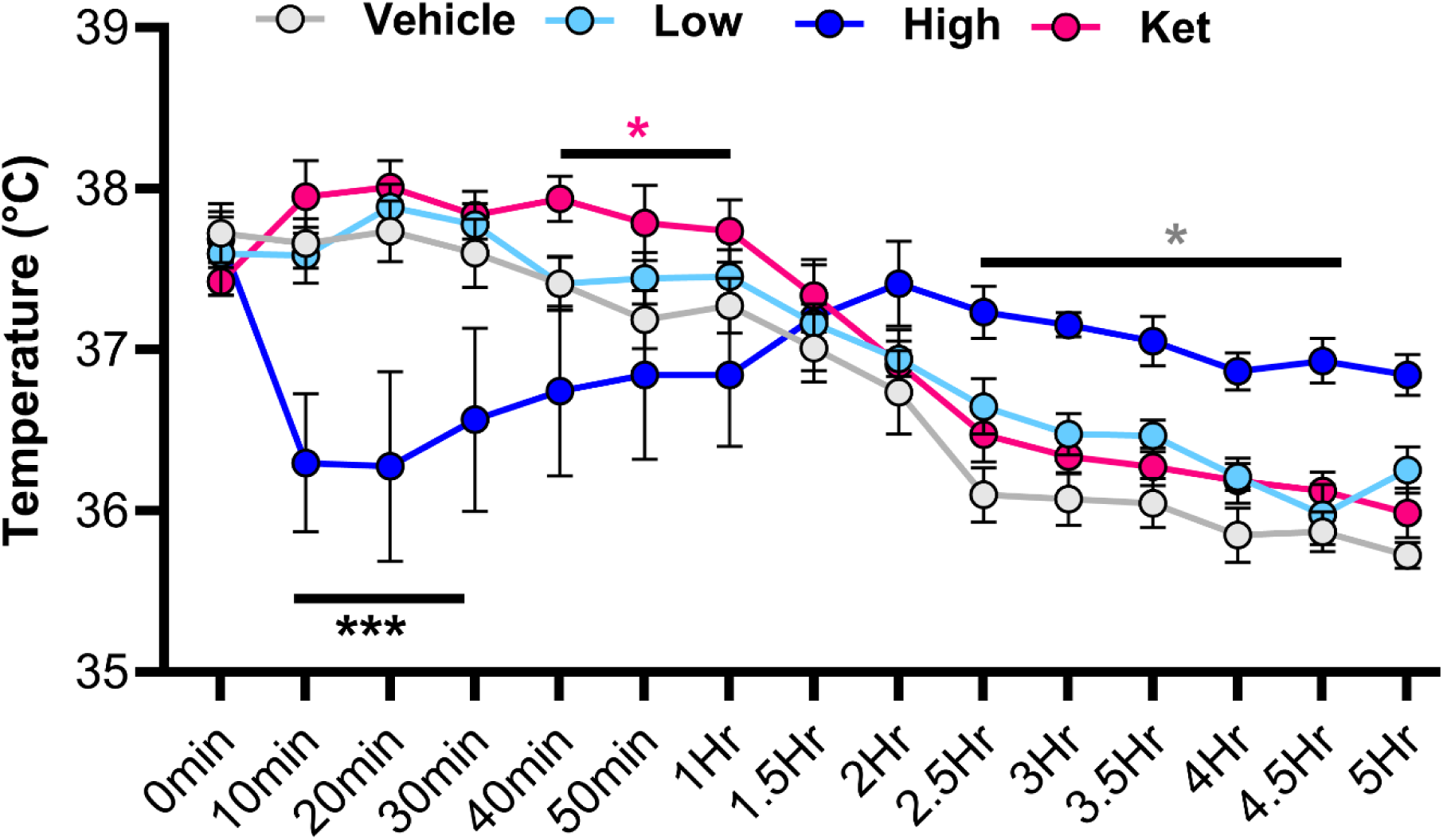
Effect of CNS4 on thermoregulation in mice. Core body temperature was monitored using a subcutaneously implanted TP-500 transponder thermometer system. Mice were briefly anesthetized during implantation and allowed to recover prior to experimentation. Baseline temperature was recorded before intraperitoneal drug administration, followed by repeated measurements up to 5 h post-dose using a noninvasive handheld wireless scanner while mice remained in their home cages. Vehicle, n=8; light blue (CNS4, 50mg/kg), n=9; blue (CNS4, 100mg/kg), n=9; ketamine, red (15mg/kg), n=8. Asterisks indicate either all (black) or a color matching group compared with 100mg/kg dose. Route, intraperitoneal. *p<0.05, ***p<0.001.

### 3.4 CNS4 does not affect fear learning, memory consolidation, but increases fear expression

During fear-conditioning training, mice exhibited a significant change in freezing behavior across time bins (F (15,450) = 33.57, P < 0.0001), indicating successful acquisition of the conditioned fear response during the tone–shock pairing session (**Figure 4A)**. In contrast, neither the main effect of treatment (F (3,30) = 0.657, P = 0.5852) nor the time vs treatment interaction (F (45,450) = 0.760, P = 0.8711) was significant. Consistent with these findings, Šídák’s multiple-comparisons tests detected no differences between treatment groups at any time point (all adjusted P > 0.05). Because drug administration occurred after completion of the training session, these results demonstrate that fear acquisition during conditioning was comparable among all groups prior to treatment.

**Figure 4.**
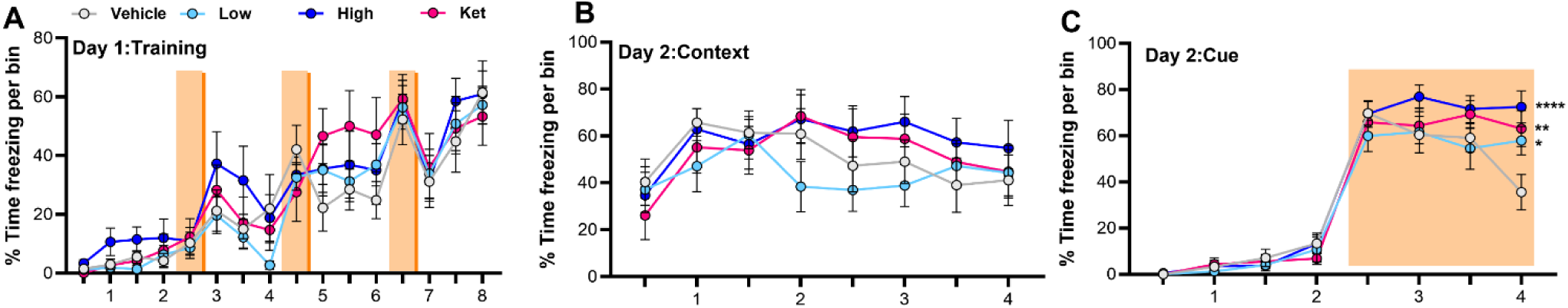
Effects of CNS4 on fear learning, contextual memory, and cued fear expression in mice. Fear conditioning was performed using a standard Pavlovian paradigm consisting of three tone-paired foot shocks (0.5 mA, 2 sec) (**A**). Each foot shock is preceded by a 30 s tone of approximately 70-dB. Contextual fear memory was assessed 24 h later by returning mice to the same chamber without tone or shock for 4 min (**B**). Cued fear memory was subsequently evaluated 1 h later in a different environment (**C**). After 2 min in silence in the new environment, the 70-dB tone was presented for a duration of 2 min. Freezing behavior was automatically recorded and quantified as an index of conditioned fear learning and memory. X axis, bin (minutes). Assigned treatments were given soon after day 1 fear conditioning training. Mice received no treatment on day 2. Vehicle, n=8; CNS4 50 mg/kg (light blue, low), n=9; CNS4 and 100 mg/kg (dark blue, high), n=9; Ketamine 15 mg/kg (purple, ket), n=8. Route, intraperitoneal. *p<0.05, **p<0.01, ****p<0.0001.

In the contextual fear memory test conducted 24 h after training, no significant effects of treatment were observed (**Figure 4B)**. Two-way repeated-measures ANOVA revealed a significant effect of time bin (F (7,210) = 3.944, P = 0.0005), indicating that freezing behavior changed over the course of the test session. However, neither the main effect of treatment (F (3,30) = 0.738, P = 0.5377) nor the time vs treatment interaction (F (21,210) = 0.910, P = 0.5784) was significant. Consistent with these findings, Šídák’s multiple-comparisons analysis identified no significant differences between any treatment groups at any time point (all adjusted P > 0.05). These results indicate that post-training administration of CNS4 (50 or 100 mg/kg) did not significantly alter the consolidation or 24-hour retention of contextual fear memory. Ketamine, likewise, did not significantly affect contextual freezing under the conditions of this experiment.

During the cued fear memory test, mice were placed in a novel context followed by tone presentation. Two-way repeated-measures ANOVA revealed a significant effect of time bin (F (7,210) = 181.6, P < 0.0001) and a significant time vs treatment interaction (F (21,210) = 1.892, P = 0.0129), while the overall treatment effect was not significant (F (3,30) = 1.708, P = 0.1866). Post hoc Šídák’s multiple-comparisons analysis showed no treatment differences during the early two bins of the test, indicating comparable baseline freezing in the novel context before or during early cue exposure (**Figure 4C)**. However, in the last two time bins, vehicle-treated mice exhibited significantly lower freezing than CNS4 50 mg/kg-treated mice (mean difference = 22.29, adjusted P = 0.0175), CNS4 100 mg/kg-treated mice (mean difference = 36.93, adjusted P < 0.0001), and ketamine-treated mice (mean difference = 27.51, adjusted P = 0.0023), **Figure 4C**. These data indicate that post-training CNS4 did not alter contextual fear retention but significantly modified the time course of cue-associated freezing, with both CNS4 doses enhancing freezing during the later phase of the cue test compared with vehicle.

### 3.5 CNS4 numerically reduced sucrose preference after footshock induced stress

Shock-threshold testing was performed prior to the sucrose preference assay to verify that treatment groups exhibited comparable sensitivity to footshock. Mixed-effects analysis revealed a significant effect of response type (F (1,29) = 66.84, P < 0.0001), with vocalization occurring at higher shock intensities than flinching (predicted means: 0.323 vs. 0.215 mA; mean difference = 0.108 mA, 95% CI: 0.081–0.135 mA), **Figure 5A**. In contrast, there was no significant effect of preassigned treatment groups (F (3,30) = 1.616, P = 0.2065) and no response type vs treatment interaction (F (3,29) = 0.561, P = 0.6451). Consistent with these findings, Tukey’s multiple-comparisons tests detected no significant differences among the vehicle, CNS4 50 mg/kg, CNS4 100 mg/kg, and ketamine-treated groups (all adjusted P > 0.15). These results indicate that all treatment groups exhibited comparable sensitivity to footshock prior to exposure to the standardized 0.5 mA stressor used before sucrose preference testing.

**Figure 5.**
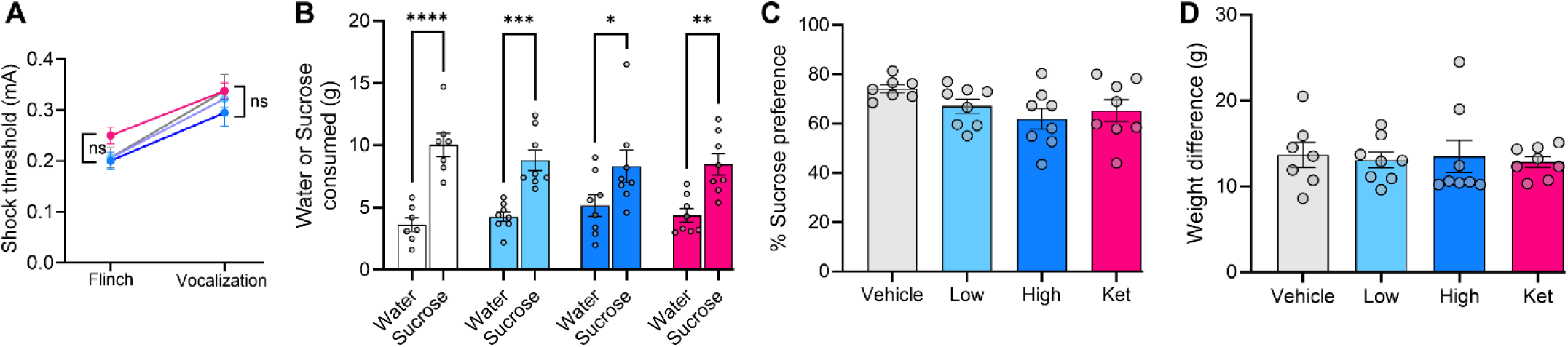
Effects of CNS4 on shock sensitivity and acute stress-induced sucrose preference in mice. Shock-threshold testing was conducted using ascending foot shocks ranging from 0.075 to 0.5 mA (1 s each) to determine behavioral responses including flinch, jump, and vocalization thresholds (**A**). Following threshold testing, mice received an acute stress-inducing 0.5 mA foot shock for 2 s prior to sucrose-preference assessment. For the sucrose-preference test, individually housed mice were provided access to water and 1% sucrose solution overnight, and bottle weights were measured before and after testing (**B**). % sucrose preference was calculated from the total liquid consumption (**C**). Difference between sucrose and water container weight is shown in **D**. Vehicle, n=8; CNS4 50 mg/kg (light blue, low), n=9; CNS4 and 100 mg/kg (dark blue, high), n=9; Ketamine 15 mg/kg (purple, ket), n=8. Route, intraperitoneal. *p<0.05, **p<0.01, ***p<0.001,****p<0.0001.

Two-way repeated-measures ANOVA revealed a significant main effect of fluid type (water versus sucrose; F (1,27) = 80.42, P < 0.0001), indicating a robust preference for sucrose across all treatment groups (**Figure 5B**). However, neither the main effect of treatment (F (3,27) = 0.081, P = 0.9697) nor the fluid type vs treatment interaction (F (3,27) = 1.734, P = 0.1837) was significant. Post hoc analyses confirmed significant sucrose preference in vehicle-treated mice (P < 0.0001), CNS4 50 mg/kg-treated mice (P = 0.0004), CNS4 100 mg/kg-treated mice (P = 0.0152), and ketamine-treated mice (P = 0.0013). However, analysis of individual sucrose preference percentages revealed mean preferences of 72.2% for vehicle, 65.3% for ketamine, 60.9% for CNS4 50 mg/kg, and 55.7% for CNS4 100 mg/kg treated mice (**Figure 5C**). Relative to the vehicle, sucrose preference was numerically reduced by approximately 9.6% in the ketamine, 15.6% in the CNS4 50 mg/kg, and 22.9% in the CNS4 100 mg/kg group. Total fluid intake did not differ significantly among the four treatment groups (one-way ANOVA, F (3, 27) = 0.0812, P = 0.9697), indicating that the observed differences in sucrose preference were not attributable to changes in overall fluid consumption (**Figure 5D**). Collectively, these findings indicate a dose-related reduction in sucrose preference following CNS4 administration, with the greatest reduction observed at 100 mg/kg.

### 3.6 Repeated administration of CNS4 does not cause changes in total body weight

Body weight was monitored on days 1, 5, 9, 12, 16, and 23 throughout the behavioral testing period, during which mice received a total of 11 assigned drug administrations (**Supplementary Figure 1**). Two-way repeated-measures ANOVA revealed a significant effect of day (F (5,150) = 61.27, P < 0.0001), reflecting normal changes in body weight over time, and a significant day vs treatment interaction (F (15,150) = 2.116, P = 0.0119). However, no significant main effect of treatment was detected (F (3,30) = 0.423, P = 0.7382). Mean body weights were similar among groups, averaging 22.83 g in vehicle, 22.98 g in CNS4 50 mg/kg, 23.20 g in the CNS4 100 mg/kg, and 23.67 g in the ketamine treated group. Consistent with the absence of a treatment effect, Tukey’s multiple-comparisons tests detected no significant differences between any treatment groups (all adjusted P > 0.72). These findings indicate that repeated administration of CNS4 did not adversely affect body weight despite producing measurable effects in several behavioral assays.

### 3.7 Single dose of CNS4 readily reverses nociceptive hypersensitivity in a rat model of neuropathic pain

A single dose of CNS4 100 mg/kg produced a marked and transient reversal of mechanical hypersensitivity in SNL rats (**Figure 6A**). Two-way repeated-measures ANOVA demonstrated a significant time vs treatment-group interaction (F (9.943, 66.29) = 9.609, P < 0.0001), as well as significant effects of time (F (3.314, 66.29) = 9.004, P < 0.0001) and treatment group (F (3,20) = 285.8, P < 0.0001). Prior to treatment, SNL rats exhibited pronounced mechanical hypersensitivity, with comparable withdrawal thresholds in the vehicle- and CNS4-treated groups (1.66 and 1.65 g, respectively). Tukey’s multiple-comparisons test demonstrated significantly greater withdrawal thresholds in CNS4-treated SNL rats than in SNL vehicle-treated rats at 30 min (adjusted P = 0.0102) and 45 min (adjusted P = 0.0413). Notably, at 30 min the withdrawal threshold in CNS4-treated SNL rats approached that of non-injured vehicle-treated controls (23.61 vs 28.32 g; adjusted P = 0.6860), indicating a substantial, albeit transient, restoration of mechanical sensitivity toward the normal range.

**Figure 6.**
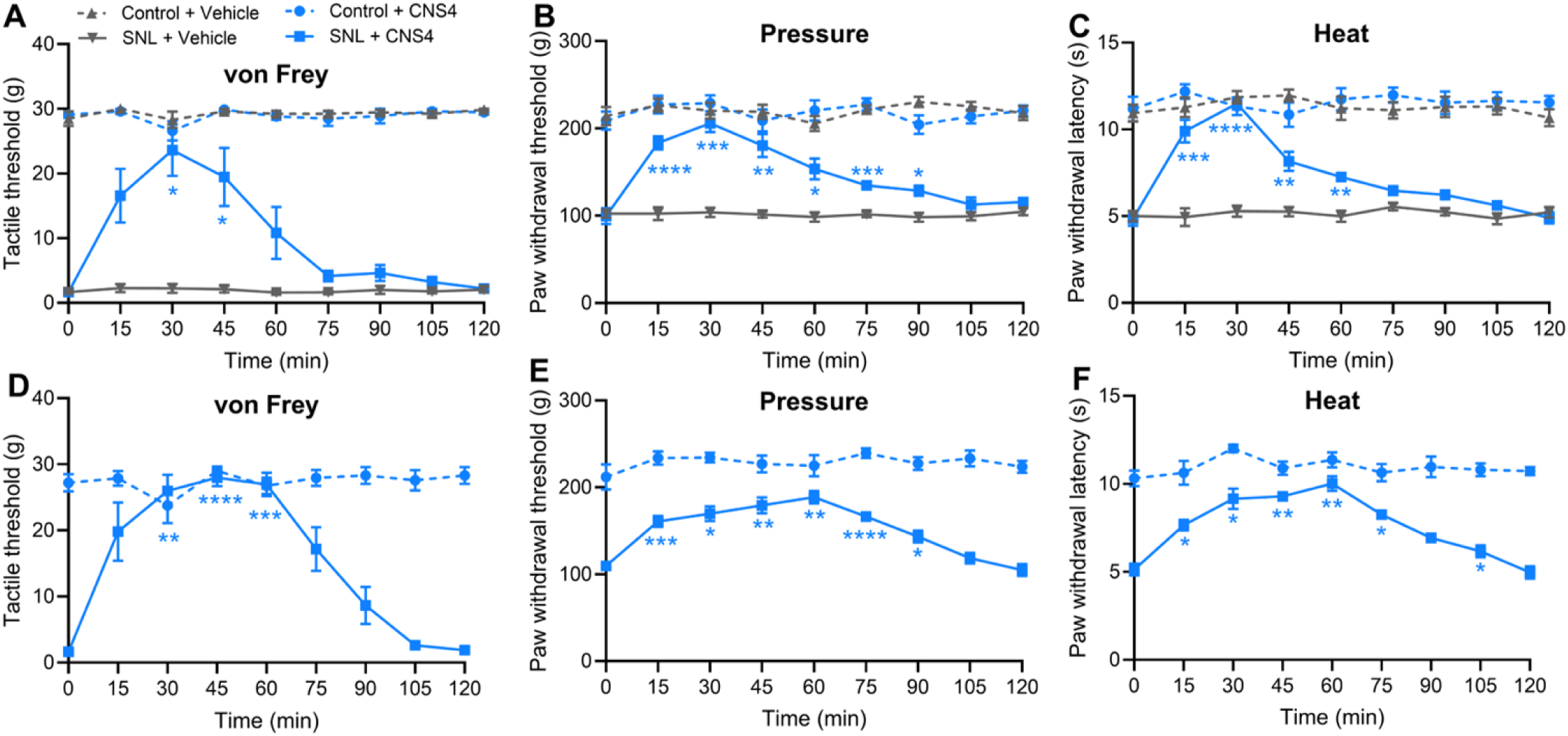
Effect of single and repeat dose CNS4 on nociceptive hypersensitivity in a rat model of neuropathic pain. The time course of the effects of a single (A–C) or three consecutive daily (D–F) intraperitoneal (IP) administrations of CNS4 (100 mg/kg) or vehicle on hind paw withdrawal responses to von Frey filaments, pressure, and heat stimuli in sham-control and SNL rats 3 weeks after surgery. Asterisks indicate statistical significance between the SNL + vehicle and SNL + CNS4 groups (A–C) or between baseline (time 0) and the indicated post-treatment time points (D–F). Statistical significance was determined by two-way repeated-measures ANOVA followed by Tukey’s multiple-comparisons test. n = 6 per group. Route, intraperitoneal. *p < 0.05, **p < 0.01, ***p < 0.001, ****p < 0.0001.

CNS4 generated a robust reversal of pressure hypersensitivity, with a significant time vs treatment-group interaction (F(15.02, 100.1) = 6.863, P < 0.0001), **Figure 6B**. Pressure withdrawal thresholds increased from 99.7 g at baseline to 183.3 g at 15 min and peaked at 206.0 g at 30 min following CNS4 treatment, compared with 104.0 g in SNL vehicle-treated rats at 30 min. CNS4 significantly increased withdrawal thresholds compared with SNL vehicle from 15 through 90 min (Tukey’s multiple-comparisons test: 15 min, P < 0.0001; 30 min, P = 0.0001; 45 min, P = 0.0055; 60 min, P = 0.0157; 75 min, P = 0.0007; 90 min, P = 0.0118).

CNS4 produced a strong reversal of thermal hypersensitivity, with a significant time vs treatment-group interaction (F(16.40, 109.4) = 7.287, P < 0.0001), **Figure 6C**. Thermal withdrawal latency increased from 4.85 s at baseline to 9.90 s at 15 min and peaked at 11.48 s at 30 min following CNS4 treatment, compared with 5.28 s in SNL vehicle treated rats at 30 min. CNS4 significantly increased withdrawal latency compared with SNL vehicle at 15 min (P = 0.0008), 30 min (P < 0.0001), 45 min (P = 0.0080), and 60 min (P = 0.0012) by Tukey’s multiple-comparisons test, with the peak response at 30 min reaching the range observed in non-injured controls.

### 3.8 Repeated administration of CNS4 maintains antinociceptive efficacy in a rat model of neuropathic pain

To determine whether antinociceptive activity was maintained following repeated administration, CNS4 100 mg/kg was administered once daily for three consecutive days, with the third dose given immediately before nociceptive testing. A separate cohort of rats were used for this set of experiments. Following repeated CNS4 administration, mechanical hypersensitivity in SNL rats was robustly reversed, with a significant time vs treatment-group interaction (F (3.354, 33.54) = 24.35, P < 0.0001), **Figure 6D**. Withdrawal thresholds increased from 1.66 g at baseline to 19.82 g at 15 min, 25.99 g at 30 min, and peaked at 27.98 g at 45 min, remaining elevated at 60 min (26.94 g). Thresholds at 30, 45, and 60 min were significantly greater than baseline (Tukey: P = 0.0028, P < 0.0001, and P = 0.0005, respectively) and were not significantly different from CNS4-treated non-SNL controls, indicating near-complete normalization of mechanical sensitivity during this period.

After repeated dose of CNS4, pressure hypersensitivity was reduced, with a significant time vs treatment-group interaction (F (3.759, 37.59) = 7.590, P = 0.0002), **Figure 6E**. Pressure withdrawal thresholds increased from 109.7 g at baseline to 160.7 g at 15 min and reached 188.7 g at 60 min, with significant increases from baseline from 15 through 90 min (Tukey: P = 0.0009, 0.0214, 0.0036, 0.0025, <0.0001, and 0.0313 at 15, 30, 45, 60, 75, and 90 min, respectively). The effect subsequently declined toward baseline by 105–120 min. Compared with the single-dose study, repeated CNS4 administration produced a more sustained effect on pressure hypersensitivity, with significantly elevated withdrawal thresholds through 90 min while retaining a similar overall magnitude of antinociceptive activity.

Similar to pressure, thermal hypersensitivity also was reduced, with a significant time vs treatment-group interaction (F (4.098, 40.98) = 7.782, P < 0.0001), **Figure 6F**. Thermal withdrawal latency increased from 5.13 s at baseline to 7.65 s at 15 min, 9.15 s at 30 min, and reached 10.02 s at 60 min; increases from baseline were significant from 15 through 75 min (Tukey: P = 0.0371, 0.0425, 0.0033, 0.0043, and 0.0141 at 15, 30, 45, 60, and 75 min, respectively). The response subsequently declined, approaching baseline by 120 min.

### 3.9 CNS4 reached systemic circulation after SC injection in client owned surgical oncology dogs

Following a single subcutaneous injection of 10mg/kg CNS4 formulation to client-owned surgical oncology dogs, plasma drug concentrations remained detectable in all animals throughout the 8-h sampling period (**Figure 7**). Mean plasma concentrations were 178.9 ± 95.2 ng/mL at 1 h, 204.2 ± 42.6 ng/mL at 4 h, and 149.8 ± 33.4 ng/mL at 8 h post-dose. Individual animal pharmacokinetic analysis yielded a mean Cmax of 235 ± 59.6 ng/mL and a mean Tmax of 3.8 ± 2.2 h, indicating relatively slow absorption following subcutaneous administration. Notably, the mean plasma concentration at 4 h exceeded that observed at 1 h (204.2 ± 42.6 vs. 178.9 ± 95.2 ng/mL), and five of the eight dogs achieved their maximum plasma concentration at 4 h or later. Together with the mean Tmax of 3.8 h, these findings suggest that absorption from the subcutaneous injection site was still ongoing during the first several hours following administration. The mean AUC0–8 was 1372 ± 264 h·ng/mL, demonstrating sustained systemic exposure throughout the postoperative observation period. Apparent elimination half-life could not be reliably estimated because plasma samples were collected only at 1, 4, and 8 h post-dose and absorption was still ongoing in most dogs during the last sampling period.

**Figure 7.**
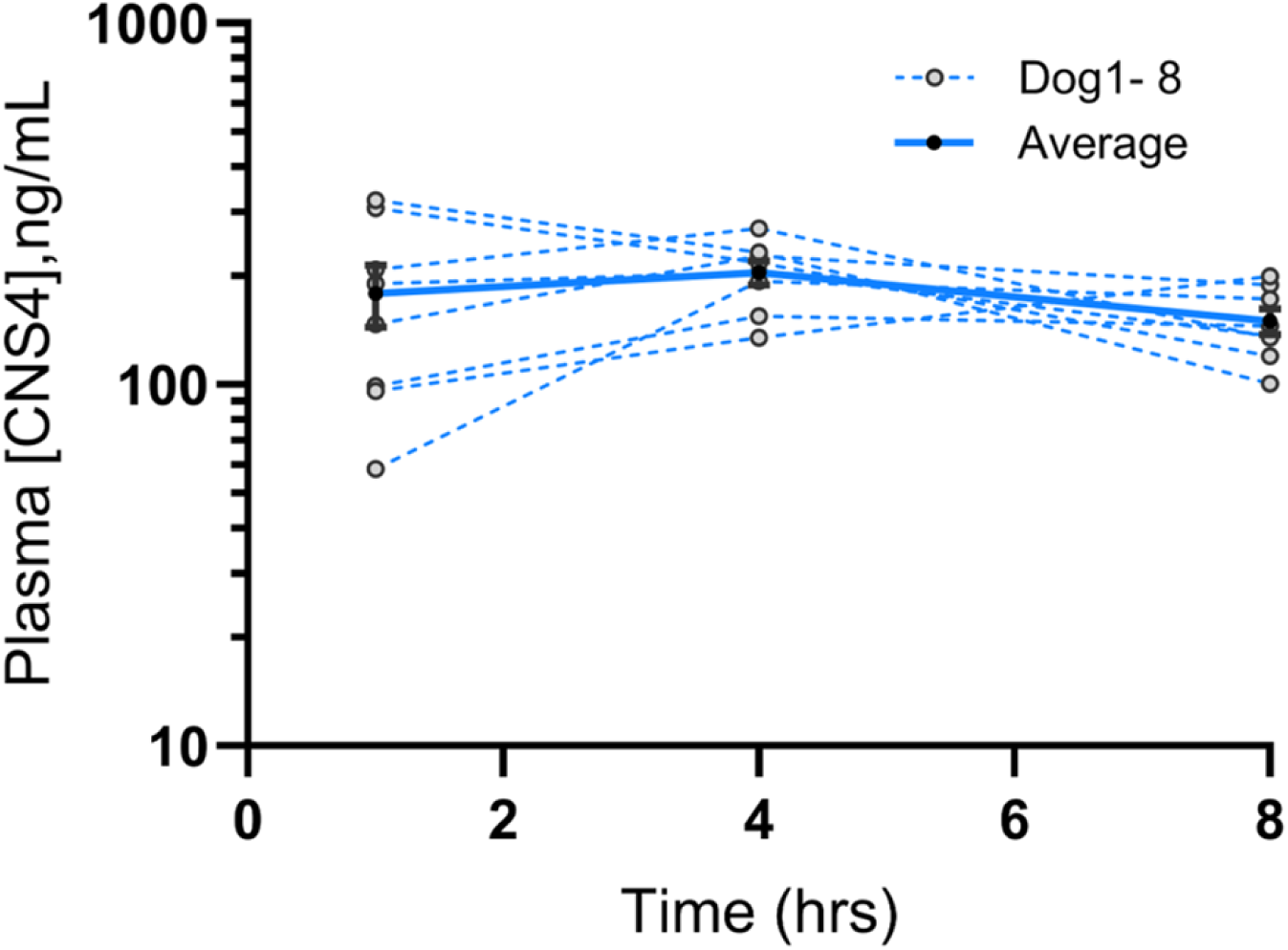
CNS4 reached systemic circulation after subcutaneous injection in client owned surgical oncology dogs. Individual (dotted line) and average (blue line) plasma concentration of CNS4 in client owned surgical oncology dogs. X-axis shows the number of hours after single subcutaneous injection of 10mg/kg CNS4. Individual and average ± sem plasma concentration of CNS4 treated dogs are plotted.

### 3.10 CNS4 supports perioperative physiologic autonomic stability

Heart rate was monitored before surgical incision, immediately after incision, and at 2, 4, and 6 h following extubation. Descriptive analysis revealed lower mean heart rates in CNS4-treated dogs during the intraoperative period, with values approximately 18.0% lower before incision (61.9 vs. 75.5 beats/min) and 17.5% lower immediately after incision (61.8 vs. 74.8 beats/min) compared with placebo-treated dogs (**Figure 8A**). Following extubation, heart rate increased progressively in both groups during recovery from anesthesia, and the difference between groups diminished. Mean heart rates at 2, 4, and 6 h post-extubation were 85.1, 89.8, and 99.4 beats/min in the CNS4 group compared with 72.7, 84.3, and 88.8 beats/min in placebo-treated dogs.

**Figure 8.**
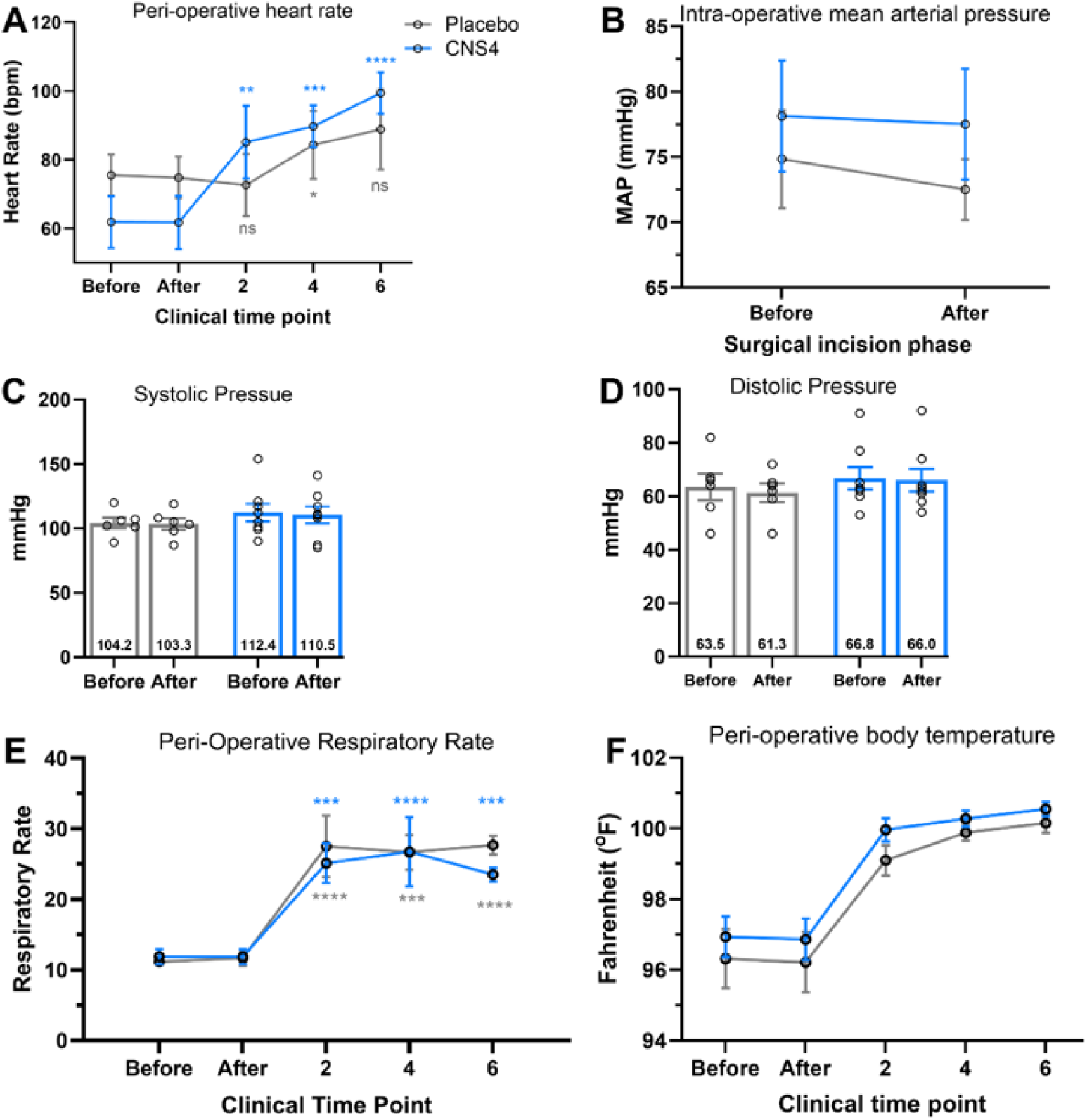
CNS4 demonstrates autonomic-stabilizing activity in dogs undergoing surgical procedures. Compared with placebo-treated dogs, CNS4-treated animals exhibited a modest reduction in heart rate (**A**) accompanied by maintenance or a slight increase in mean arterial pressure (**B**) which is derived from systolic (**C**) and diastolic (**D**) blood pressure. No sign of respiratory depression (**E**) and mild elevation in body temperature (**F**) throughout the peri-operative phase demonstrate cardiovascular stability. Before and after on the x-axis indicate measurement time points before and after surgical incision. 2, 4 and 6 hours are post extubation time points. Heart rate and respiratory rate were analyzed using two-way repeated-measures ANOVA and Šídák’s multiple-comparisons test to find the differences between before incision and post extubation time points within the same treatment group. Color matching asterisks indicate the significant level compared to before incision and respective post extubation time points. CNS4, n=8; Blue, placebo, n=6. Route: subcutaneous. *p<0.05. **p<0.01, ***p<0.001, ****p<0.0001.

Mean arterial pressure (MAP) was recorded immediately before and after surgical incision. Because CNS4 was administered approximately 90 minutes before the pre-incision measurements, the values obtained before incision reflect the early perioperative effects of CNS4 exposure. CNS4 treated dogs exhibited numerically higher MAP values than placebo-treated dogs both before incision (78.1 vs. 74.8 mmHg; +4.4%) and after incision (77.5 vs. 72.5 mmHg; +6.9%), **Figure 8B**. Examination of the blood pressure components revealed that CNS4 treated dogs also maintained numerically higher systolic and diastolic pressures throughout the peri-incisional period. Mean systolic blood pressure was 112.4 mmHg before incision and 110.5 mmHg after incision in the CNS4 group, compared with 104.2 and 103.3 mmHg, respectively, in placebo-treated dogs (+7.9% before and +7.0% after incision), **Figure 8C**. Likewise, mean diastolic pressure was 66.8 mmHg before incision and 66.0 mmHg after incision in CNS4-treated dogs, compared with 63.5 and 61.3 mmHg in placebo-treated dogs (+5.2% before and +7.7% after incision) **Figure 8D**. MAP remained stable across the peri-incisional period in both groups, with only a 0.8% decrease in the CNS4 group compared with a 3.1% decrease in placebo-treated dogs. Notably, preservation of MAP in CNS4-treated dogs appeared to be driven primarily by maintenance of diastolic pressure, which declined by only 1.2% in the CNS4 group compared with 3.5% in placebo-treated dogs. Further, CNS4-treated dogs required approximately 8% less propofol for anesthetic induction than placebo-treated dogs (2.60 ± 0.57 vs. 2.83 ± 0.58 mg/kg). Despite the heterogeneity of the surgical procedures, propofol requirements were numerically lower in the CNS4-treated group.

Respiratory rate was monitored before surgical incision, immediately after incision, and at 2, 4, and 6 h following extubation. Mean respiratory rates were similar between CNS4 treated and placebo treated dogs during the intraoperative period (11.9 vs. 11.2 breaths/min before incision and 11.9 vs. 11.7 breaths/min after incision), **Figure 8E**. Following extubation, respiratory rates increased in both groups as expected during recovery from anesthesia, reaching mean values of 25.1, 26.8, and 23.5 breaths/min in the CNS4 group and 27.5, 26.7, and 27.7 breaths/min in the placebo group at 2, 4, and 6 h post-extubation, respectively. These observations indicate that CNS4 administration was not associated with respiratory depression or impaired postoperative ventilatory recovery. However, interpretation of intraoperative respiratory rates is complicated by the use of anesthetic medications and respiratory support, including mechanical ventilation when clinically indicated, which may have influenced respiratory rate and limited assessment of spontaneous hypoventilation during surgery.

### 3.11 Potential contribution of CNS4 to perioperative thermoregulatory homeostasis

Body temperature was monitored before surgical incision, immediately after incision, and at 2, 4, and 6 h following extubation. As expected, both groups exhibited mild perioperative hypothermia during anesthesia, with mean temperatures of 96.9°F and 96.9°F in the CNS4-treated group and 96.3°F and 96.2°F in placebo-treated dogs before and immediately after incision, respectively (**Figure 8F**). Following extubation, body temperature progressively returned toward the normal physiologic range in both groups. Notably, CNS4-treated dogs maintained numerically higher temperatures throughout the perioperative period, with mean values approximately 0.4–0.9°F greater than placebo-treated dogs at all measured time points. Since body temperature is a highly sensitive physiologic parameter during anesthesia and surgical recovery, a mild but sustained increase is remarkable.

## 4. Discussion

The development of the CNS4·HCl salt is a major advancement in this drug development study on the chemistry, manufacturing, and controls (CMC) aspect. Although free base was sufficient for proof-of-concept efficacy, pharmacokinetic, and safety studies (4, 5); its formulation characteristics were less compatible with the requirements of long-term preclinical development and eventual clinical use. Conversion to the hydrochloride salt substantially improved solubility in physiologically relevant vehicle, 20% HPβCD in water at concentrations up to 30 mg/mL. Thus, while the free base established the pharmacological activity of CNS4, the hydrochloride salt provides a more practical and clinically translatable form for future GLP toxicology and IND-enabling studies.

NMDAR antagonists, particularly ketamine and its parent compound phencyclidine, produce psychomotor stimulation, reinforcing effects, and abuse liability (18, 19). However, mice that received CNS4 for four consecutive days did not exhibit conditioned place preference (**Figure 1**). This finding suggests that CNS4, which is chemically distinct from NMDAR channel blockers, may also act through a distinct mechanism in vivo. Cocaine was chosen as a positive control for the CPP assay, as this is more frequently used in mice compared to ketamine, which at doses needed to cause addiction also can cause sedation, a confounding factor for CPP assay (20, 21). However, cocaine caused only numerical increase in CPP. This is probably due to variations in individual animals and small sample size, n=8. Anecdotally, one mouse in cocaine group had a reversal learning of saline vs drug chamber (**Figure 1B&C**) but this was not excluded from the analysis.

The behavioral profile of CNS4 differed substantially from that typically reported for NMDAR channel blockers. NMDAR channel blockers are known to cause hyperlocomotion in rodents (22, 23). In contrast to this, high dose CNS4 markedly reduced spontaneous locomotor activity, whereas the low dose had little effect, indicating dose-dependent modulation of neural circuits involved in arousal and exploratory behavior (**Figure 2A**). However, interestingly, despite reduction in locomotion, CNS4 did not impair motor coordination or motor learning as assessed by rotarod performance, nor did it interfere with fear acquisition, memory consolidation, or context- and cue-dependent fear recall (**Figure 2B &4**). Indeed, the highest level of fear memory recollection at cue test (p<0.001 compared to vehicle) was observed with high dose CNS4 (**Figure 4C**). These findings suggest that CNS4 does not produce generalized motor suppression or cognitive impairment but instead selectively reduces spontaneous locomotion while preserving motor and cognitive function. Similar cognitive maintenance was also reported in a previous fear conditioning study in which CNS4 was injected before the fear conditioning training (4). Such a profile is consistent with the NMDAR subtype-dependent pharmacology of CNS4, which modulates GluN2-containing NMDARs in a manner distinct from nonselective channel-blocking antagonists, potentially allowing regulation of arousal-related glutamatergic circuits without disrupting the NMDAR-dependent synaptic plasticity required for motor learning and memory formation.

NMDAR mediated glutamatergic signaling contributes to central thermoregulation, including within the hypothalamic medial preoptic area, a key region involved in integration and control of body temperature (24, 25). Pharmacological inhibition of NMDAR signaling has been associated with impaired thermoregulation and hypothermia, and high dose of ketamine is known to reduce body temperature in rodents (26, 27). In the present study, high-dose CNS4 produced a biphasic thermoregulatory response characterized by a transient early decrease in body temperature followed by a sustained elevation above vehicle values during the later observation period (**Figure 3**). Importantly, these temperature changes remained within the normal physiological range and did not constitute overt hypo- or hyperthermia. A similar trend was observed in client-owned surgical oncology dogs, where CNS4-treated animals maintained slightly higher body temperatures during the entire perioperative period compared with placebo-treated dogs (**Figure 8**). Given the role of NMDAR-dependent glutamatergic signaling in hypothalamic thermoregulatory pathways, these findings suggest that CNS4 modulates rather than suppressing thermoregulatory network activity. This profile is consistent with the previously reported mechanism of CNS4 as a subtype-dependent modulator that might preserve physiological NMDAR function while altering network excitability (1, 3).

A reduction in sucrose preference observed following acute stress exposure suggests that CNS4 modulates neural circuits involved in reward processing and stress responsiveness (**Figure 5**). NMDARs play a central role in synaptic plasticity within corticolimbic networks, including the prefrontal cortex, hippocampus, nucleus accumbens, and amygdala, which collectively regulate motivation, reward valuation, and affective behavior (28). Dysregulation of glutamatergic signaling within these circuits has long been implicated in depression and stress-related disorders (29–31). Interestingly, both CNS4 (22.9%) and ketamine (9.6%) reduced stress-induced sucrose preference to a similar extent, despite their markedly different pharmacological mechanisms. Ketamine, widely regarded as one of the most important advances in biological psychiatry in the last 50 years because of its rapid antidepressant effects, is thought to act through transient NMDAR blockade, leading to downstream enhancement of synaptic plasticity and neurotrophic signaling (16). In contrast to ketamine’s mechanism of action (MOA), CNS4 is a biased NMDAR modulator that likely preserves receptor function in fluctuating glutamate concentration (1, 3). Despite the differences in the currently known MOA, CNS4 reduced sucrose preference numerically better than ketamine suggesting that beneficial modulation of NMDAR signaling may be achieved without channel blockade. Although reduced sucrose preference in itself is not definitive indicator of antidepressant activity, when considered together with the absence of cognitive impairment, preservation of fear learning and memory, reduced behavioral activation, and the lack of abuse liability observed in the conditioned place preference assay, these findings support the hypothesis that CNS4 engages stress-responsive glutamatergic circuits through a mechanism distinct from ketamine while retaining potentially favorable neuropsychiatric drug like properties.

Neuropathic pain is associated with maladaptive glutamatergic synaptic plasticity within the spinal dorsal horn, where primary nociceptive afferents terminate and relay sensory information to ascending pain pathways (13, 32, 33). GluN1/2A, GluN1/2B, GluN1/2A/2B, and GluN1/2D subtypes of NMDARs play a central role in this process by mediating activity-dependent synaptic strengthening and central sensitization (34, 35). The contribution of NMDARs to both physiological and pathological pain signaling remains complex and context dependent (13, 33). Thus, CNS4’s concentration-dependent and GluN2 subtype-dependent modulation, enhancing receptor function under low glutamate conditions while reducing hyperactivation at higher glutamate concentrations, may be particularly advantageous for neuropathic pain, in which extracellular glutamate levels are elevated within spinal nociceptive pathways. Results obtained from the rat SNL model (**Figure 6**) support the hypothesis that CNS4 attenuates neuropathic pain through modulation of NMDAR dependent central sensitization through dorsal horn nociceptive circuits. The ability of CNS4 to restore withdrawal thresholds toward normal values without altering control animal sensitivity to nociception is a significant therapeutic advantage (**Figure 6A&D**). Such a mechanism could potentially preserve physiological synaptic plasticity and cognitive function while selectively reducing pathological glutamatergic signaling that drives chronic pain. Further, retention of robust antinociceptive activity following three consecutive daily doses indicates that CNS4 efficacy was maintained during the repeated-dose regimen (**Figure 6D-F**), with no apparent evidence of rapid tolerance development. This finding is noteworthy because rapid loss of antinociceptive efficacy can occur with repeated administration of some analgesics, particularly opioids such as morphine (36). Although near-complete reversal of neuropathic hypersensitivity has been previously reported with potent opioid/NOP agonists, cannabinoids, gabapentinoids, and TRPV1-directed interventions (37–40), the broad efficacy of CNS4 across tactile, pressure, and thermal modalities is remarkable for a mechanistically distinct NMDAR modulator. To our knowledge, no previously described NMDAR modulator with a comparable pharmacological profile has demonstrated this degree of efficacy across multiple sensory modalities in rat neuropathic pain model.

NMDARs are widely expressed throughout motor and autonomic regulatory networks (41–43). Within cortical and subcortical motor circuits, NMDAR-mediated neurotransmission contributes to neuronal excitability, activity-dependent synaptic plasticity, motor learning, and regulation of locomotor behavior (44, 45). In parallel, NMDAR activation within the rostral ventrolateral medulla (RVLM) regulates sympathetic vasomotor outflow and arterial pressure, whereas NMDAR signaling within the dorsal motor nucleus of the vagus (DMV) contributes to parasympathetic control of cardiac and visceral function (46–50). Consequently, modulation of NMDAR activity by CNS4 may influence regulation of heart rate, vascular tone, arterial blood pressure, respiratory drive, and thermoregulation. Results from the perioperative study in client-owned oncology dogs with advanced, naturally occurring cancer suggest that CNS4 may influence autonomic regulatory responses under the physiological stress of anesthesia, surgery, and postoperative recovery (**Figure 8**). CNS4-treated dogs exhibited lower peri-incisional heart rate while maintaining numerically higher MAP, systolic pressure, and diastolic pressure compared with placebo-treated mice. Preservation of diastolic pressure is particularly notable because diastolic pressure is a major determinant of MAP and reflects maintenance of peripheral vascular resistance. Thus, CNS4-treated dogs appeared capable of maintaining arterial pressure despite lower heart rates, a pattern suggestive of improved cardiovascular efficiency rather than generalized sympathetic activation. Consistent with this interpretation, CNS4-treated dogs also demonstrated comparable respiratory rates and slightly higher body temperatures during recovery, both of which are sensitive physiologic indicators of autonomic function and recovery from anesthesia. Overall, these findings are particularly notable given the heterogeneous clinical population, in which individual dogs differed in underlying cancer, disease burden, and physiological status, yet CNS4-treated animals maintained cardiopulmonary and autonomic stability during the substantial physiological stress of anesthesia and surgery.

### Limitation

Postoperative pain scores and the analgesic burden score (ABS) were evaluated to explore the potential analgesic effect of CNS4 in dogs. However, pain score itself was not suitable as a direct efficacy endpoint because the clinical protocol required administration of rescue analgesics whenever a dog reached a predefined pain threshold (score ≥2), thereby preventing sustained differences in pain scores between treatment groups. Consequently, ABS, which reflects the amount and frequency of additional analgesic medication required to maintain adequate pain control, may provide a more informative measure of potential analgesic benefit. CNS4-treated dogs showed a numerically lower ABS compared with placebo-treated dogs, providing a preliminary signal of an opioid/analgesic-sparing effect. Nevertheless, this interpretation is limited by substantial clinical heterogeneity, including differences in patient weight, tumor type and size, surgical complexity and duration, tissue trauma, and perioperative anesthetic requirements. Therefore, these observations could not be included in this manuscript. Thus, this aspect of the work warrants further investigation in a larger, prospectively controlled study.

## 5. Conclusions

The maintenance of cardiovascular parameters and body temperature, respiratory recovery, and reduced analgesic requirement in dogs are consistent with CNS4 supporting physiologic stability during perioperative stress. These findings complement the antinociceptive effects observed in rats, in which CNS4 reduced responses to thermal and mechanical nociceptive stimuli, supporting its pharmacologically relevant analgesic activity. In mice, CNS4 maintained body temperature and markedly reduced spontaneous locomotor activity while no detectable impairment of motor learning. Together, these findings across rat, mouse, and canine models suggest that concentration-dependent and subtype-dependent modulation of NMDARs by CNS4 may provide analgesic activity while preserving motor and autonomic functions. These properties support further translational development of CNS4 for neuropathic pain and potentially neuropsychiatric disorders.

## 6. Ethics declaration

All animal experiments were conducted in accordance with applicable institutional guidelines and were approved by the Institutional Animal Care and Use Committees (IACUCs) of the respective institutions where the studies were performed. Studies involving client-owned dogs were conducted under the approved Virginia Tech IACUC/clinical research protocol, and written informed consent was obtained from the owners prior to enrollment.

## 7. CRediT authorship contribution statement

Joanne Tuohy: Investigation, Methodology, Funding acquisition, Writing – review & editing. Toshitsugu Ishihara: Investigation, Methodology, Writing – review & editing.

Sudarshana Govindasamy: Investigation, Methodology

Ramu Anandakrishnan: Formal analysis, Writing – review & editing Jennifer Davis: Investigation, Methodology, Formal analysis

Fiona E. Harrison: Investigation, Methodology, Formal analysis, Writing – review & editing Yuying Huang: Investigation, Methodology

Shao-Rui Chen: Investigation, Methodology

Hui-Lin Pan: Investigation, Methodology, Formal analysis, Writing – review & editing

Blaise M. Costa: Conceptualization, Investigation, Methodology, Project administration, Supervision, Funding acquisition, Writing – original draft, Writing – review & editing

## 8. Declaration of generative AI and AI-assisted technologies in the manuscript preparation process

During the preparation of this work the authors used ChatGPT to generate a graphical figure. After using this tool/service, the authors reviewed and edited the content as needed and take full responsibility for the content of the published article.

## 9. Declaration of competing interest

BMC is the inventor on U.S. Patent No. 12,485,115, “Biased NMDA Receptor Modulators and Uses Thereof,” which covers CNS4 and related compounds. BMC is also the founder and CEO of Clab LLC. The other authors declare no competing financial interests or personal relationships that could have influenced the work reported in this paper.

## 10. Acknowledgments

This work was supported by the FY2025 and FY2026 Edward Via College of Osteopathic Medicine (VCOM)–Virginia Tech One Health Research Grant funded to JH and BMC.

## 11. Appendix A. Supporting information

Supplementary data associated with this article can be found in the online version.

## 12. Data availability

Data will be made available on request.

---

**Supplementary Figure 1.**
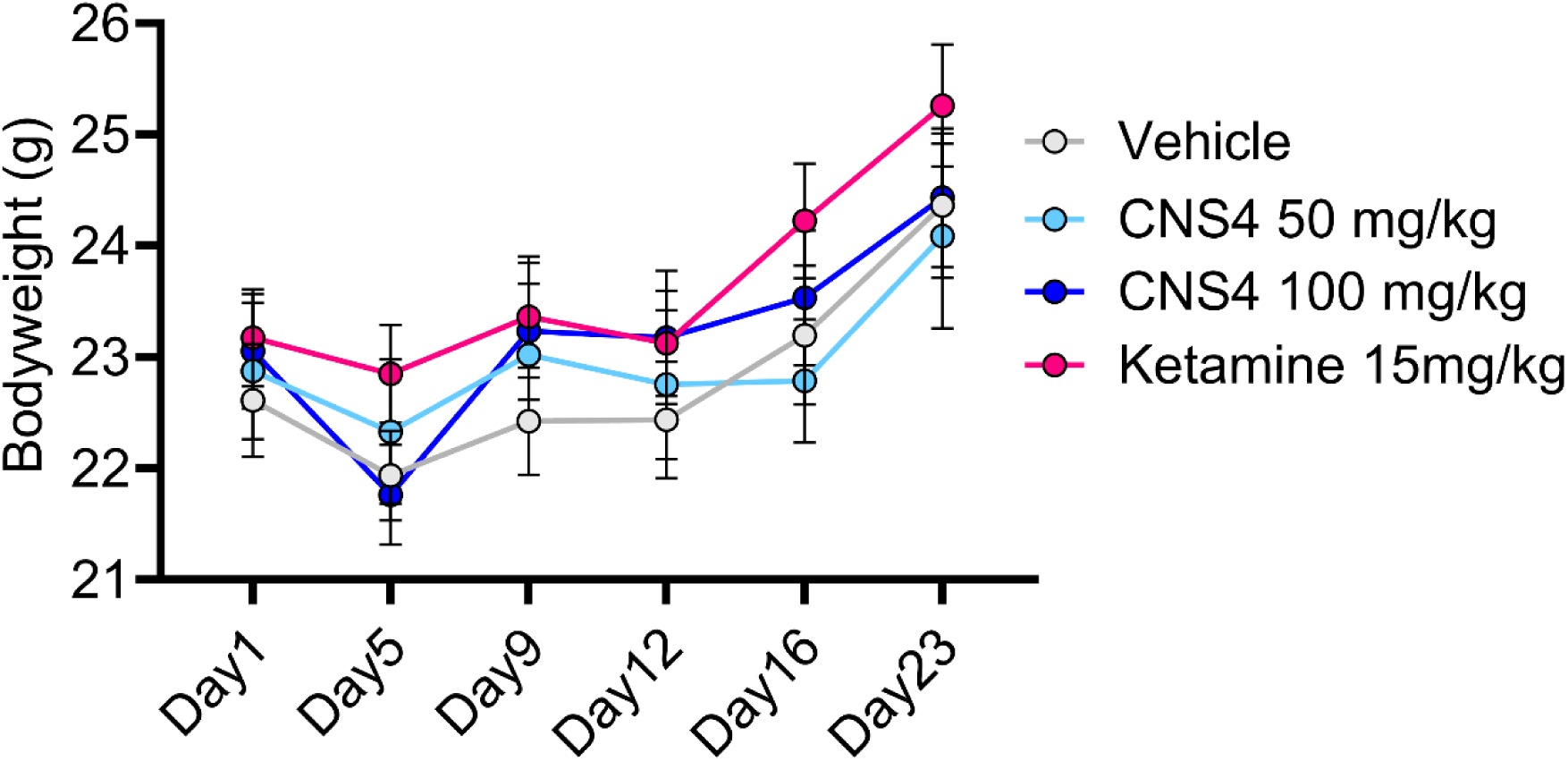
Effect of repeated CNS4 treatment on body weight in mice. Body weight was monitored over the 23-day study period, during which mice received their assigned treatment (vehicle, CNS4 at 50 or 100 mg/kg, or ketamine at 15 mg/kg) a total of 11 times. Body weight changed significantly over time (P < 0.0001), with a significant time vs treatment interaction (P = 0.0119), but no significant overall treatment effect (P = 0.7382; two-way repeated-measures ANOVA). Overall, repeated CNS4 administration did not produce a sustained reduction in body weight compared with vehicle. Data are presented as mean ± SEM.

